# Temporal development of *Drosophila* embryos is highly robust across a wide temperature range

**DOI:** 10.1101/309724

**Authors:** Jeronica Chong, Christopher Amourda, Timothy E. Saunders

**Affiliations:** Mechanobiology Institute, National University of Singapore, Singapore; Department of Biological Sciences, National University of Singapore, Singapore; Institute of Molecular and Cell Biology, A*Star, Proteos, Singapore

**Author notes:** These authors contributed equally to this paper (Jeronica Chong, Christopher Amourda).

## Abstract

Development is a precisely coordinated process in both space and time. Spatial precision has been quantified in a number of developmental systems, and, for example, such data has contributed significantly to our understanding of morphogen gradient interpretation. However, comparatively little quantitative analysis has been performed on timing and temporal coordination during development. Here, we use *Drosophila* to explore the temporal robustness of embryonic development within physiologically normal temperatures. We find that development is temporally very precise across a wide range of temperatures in all three *Drosophila* species investigated. However, we find temperature dependence in the heterochronicity. A simple model incorporating history-dependence can explain the developmental temporal trajectories. Interestingly, the history-dependence is temperature specific with either effective negative or positive feedback at different temperatures. We also find that embryos are surprisingly robust to shifting temperatures during embryogenesis. We further identify differences between tropical and temperate species that are suggestive of different potential mechanisms regulating temporal development depending on the local environment. Overall, our data shows that *Drosophila* embryonic development is temporally robust across a wide range of temperatures but there are species specific differences.

## Introduction

Multicellular organism development is characterised by the ability to complete morphogenesis with little variation between individuals. In particular, quantitative experiments on patterning processes early in embryogenesis have shed important light into this level of reproducibility (1-4). During early embryonic development, coarse gradients are subsequently refined to reach a pattern resolved at the single-cell level (5). Remarkably, the spatial precision of patterning often remains unaffected in the face of environmental fluctuations within typical physiological ranges. For example, the fly wing vein patterning operates at the physical limit (i.e. at the single-cell level) and that this limit is robust to a wide temperature range (6). Whilst a lot of attention has been brought to the level of spatial precision, developmental time precision is comparatively poorly studied. In the *Drosophila* embryo, the time to hatching roughly doubles upon a temperature change from 21°C to 16°C. Therefore, exploring temporal reproducibility is essential to gain insights into the organism response to environmental changes.

Endotherm animals maintain relatively constant body temperature, within a degree or two from their optimal temperature. Past studies have shown that a combination of metabolic and behavioural responses sustains body temperature (7). In contrast, ectotherm animals are unable to regulate their body temperature and, therefore, need to physiologically adapt to fluctuating conditions (7). Similarly, proper behavioural response is pivotal in maintaining ectotherm organisms in adequate living temperature (8,9). For example, the genes Painless and Pyrexia are critical for high-temperature nociception in *Drosophila* larvae (10-12). In their absence, larvae exhibit latencies in sensing and moving to colder temperatures. The porcelain crab, *Petrolisthes,* remains under stones during low tides when the temperature may raise over 20°C in 6 hours (13). Kenyan chameleons (*Chameleo dilepis* and *Chameleo jacksonii*) alter their skin colouration to a darker tone in order to effectively absorb early morning sun, allowing them to reach their optimal temperature faster (14). Furthermore, sex determination of several species of reptiles including crocodiles and most turtles are particularly sensitive to temperature (15). A hotter environment is correlated with increased levels of aromatase, an enzyme converting androgens to estrogens. Therefore, hot temperatures direct gonads differentiation to the female fate whilst colder temperature induces male fate. Such behavioural responses are instrumental to maintain organism viability in varying environmental conditions. How temperature affects developmental time has also been widely studied (16-18). Cross-species analysis has even found evidence for a “biological clock” that links developmental time with temperature and body size (19).

The ability to respond to environmental changes is unequal throughout the life cycle of any ectotherm animal. Whilst *Drosophila* larvae and adults clearly exhibit acute behavioural responses to environmental changes (8), the embryonic stage is unable to do so and is, therefore, vulnerable to perturbations (20). It has been hypothesised that female flies can improve offspring fitness by depositing eggs in thermally favourable locations (21), though this appears unlikely as the temperature at a given time does not reflect the future temperature. In particular, the *Drosophila* embryo typically experiences at least one day/night cycle with corresponding temperature changes. Given the apparent vulnerability of *Drosophila* embryos, it is important to understand whether embryos exhibit significant changes in developmental time precision at certain temperatures and how they respond to varying environments.

Despite the importance of temporal adaption to development, the quantification of developmental time is considerably less comprehensive than analogous studies of spatial precision. Here, we began by asking: is *Drosophila* embryonic development temporally robust at different temperatures? By robust, we mean that the heterochronicity (fluctuations in developmental time) between embryos under equal conditions are on the order of a few percent (as a percentage of mean time), which is comparable to the robust spatial boundaries defined in the *Drosophila* embryo. We find that embryonic development of three *Drosophila* species (*D. melanogaster, D. simulans and D. virilis*) are all temporally robust across a broad range of temperatures, with relative errors comparable to the relative error in spatial positioning of many gene boundaries. Having quantified the temporal robustness of *Drosophila* embryonic development we then asked: (i) is development temporally robust to temperature variability; and (ii) does the temporal robustness display temperature-dependent behaviour? A combination of temperature shifts and pulse experiments in *D. melanogaster* reveals that temporal robustness is sensitive to temperature fluctuations in early embryogenesis, but the temporal error does not increase substantially. Further, we find that the statistical properties of the heterochronicity are temperature dependent, with embryogenesis most temporally robust around intermediate (19°C-23°C) temperatures. We are able to explain this observation through a simple model that incorporates history-dependence of the temporal trajectories through development. Finally, we discuss differences in the temporal robustness between the different *Drosophila* species, and our work highlights potential differences in how temperate and tropical species temporal adapt to temperature changes.

## Materials and methods

### Fly stocks

We used in-bred *Drosophila melanogaster (D. melanogaster)* OregonR, *Drosophila simulans (D.simulans)* Rakujuen and *Drosophila virilis (D. virilis)* viri-HUE lines to minimize genetic diversity in our samples. Flies were maintained with standard fly food containing cornflour, dextrose, yeast brewers, Bacto Agar and 10% Nipagin. Flies were kept at 25°C through all life cycles. Prior to imaging, flies were caged and kept at 25°C. Flies were allowed to lay on an apple juice agar plate (Agar, Sucrose and Apple juice) where the embryos were collected. Only imaged embryos were subjected to different temperatures.

### Sample preparation

Twenty non-dechorionated embryos were aligned on an apple juice agar plate (figure 1*a*). Embryos were selected at the blastoderm stage and allowed to develop at a precise temperature (in nearly all experiments the temperature fluctuations δT were very small compared to the temperature (δT/T<0.5%, figure 1*b*) until hatching in Halocarbon oil 27 to visualise developmental stages. The embryos were imaged on a Nikon SMZ18 stereomicroscope appended with a Julabo GmbH temperature control device. The temporal resolution was 2 minutes. All experiments were performed on the same microscope setups with identical illumination strength. At constant temperatures, temperature shift and fluctuation experiments, embryo survival rate (defined by whether larvae hatched) was >70% (figure S1*a-d*). Most experiments were repeated at least three times, with a minimum of 35 embryos in each temperature batch (and greater than 70 for most temperatures). We repeated experiments at 16, 21, and 25°C in *D. melanogaster* over a year apart to check that our results were robust to experimental drift, such as different batches of food and multiple generations later.

**Figure 1:**
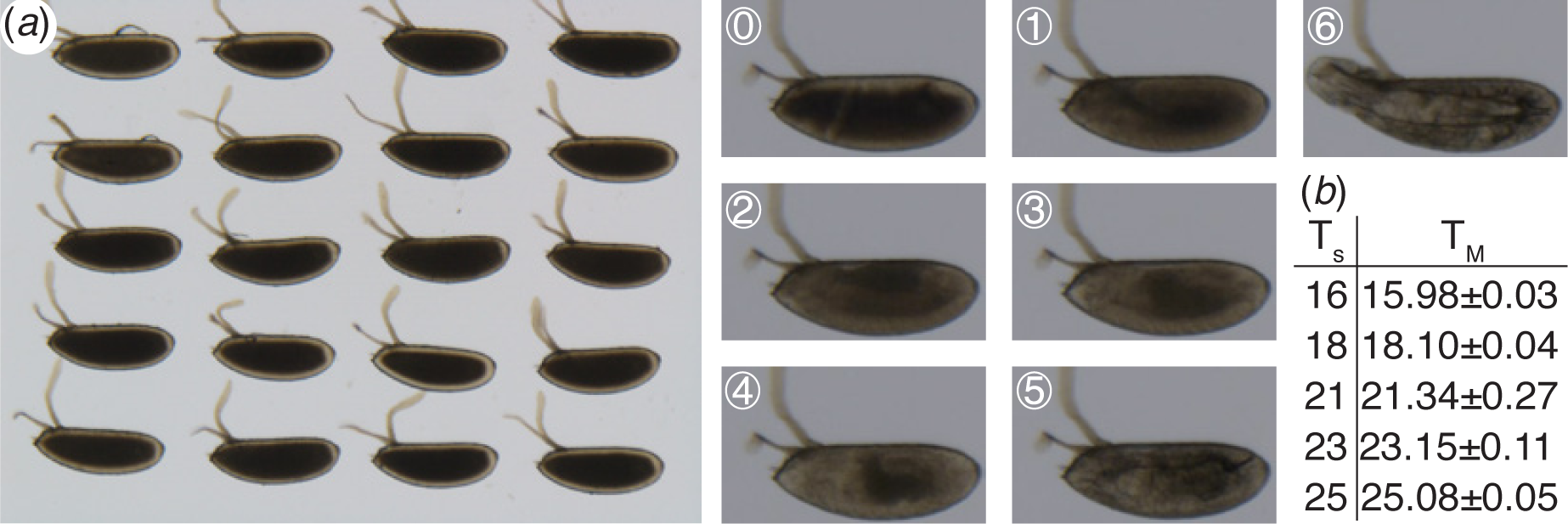
Experimental setup. (*a*) Left: Image of 20 *D. melanogaster* embryos in the microscope setup. Right: Highlighting the different landmarks used in the paper to analyse temporal development, see also Materials and Methods. (*b*) Measured temperatures compared with the temperature set and corresponding standard deviation.

### Image analysis

The developmental time was scored based on seven developmental landmarks (figure 1*a* and Table 1). We used the cephalic furrow formation as time 0 to set each embryo to a common start time. Further, we scored the time of germband retraction, head involution, midgut broadening, muscle contraction, trachea filling and hatching to span the entire embryogenesis at regular interval. The cephalic furrow formation is a transient process and constitutes the first event of gastrulation. Invagination of cells occurs on the lateral side of the embryo at about 65% of the embryo length (from the posterior of the embryo). The germband retraction stage is identified when the germ band shortens at the dorsal side of the embryo. Head involution occurs midway through embryogenesis, due to internalization of the ectodermal tissue and rearrangement of cells. The midgut broadening landmark is identified by the formation of a triangular-shape laterally. Muscular movement is characterised by uncontrolled twitching of muscles. Before hatching, the tracheal tree is filled with air and is visible due to the rapid darkening of the trachea. Last, we scored the time when larvae hatch from the embryonic case. Landmark identification was done visually and by the same experimenter for all movies. To minimise potential bias, the experimenters making the measurements and analysing the data were different.

**Table 1.** Developmental landmarks used to time *Drosophila* embryonic development.

| Label | Stage | Description |
| --- | --- | --- |
| 0 | Cephalic furrow formation | Ingression of cells around 30% embryo length, starting with cells on the lateral sides of the embryo |
| 1 | Germband retraction | Retraction of germband from dorsal side of embryo, with embryo transitioning from parasegmental to segmental division |
| 2 | Head involution | Internalisation and rearrangements of head segments |
| 3 | Midgut broadening | Expansion of gut structures |
| 4 | Muscle contractions | Uncontrolled twitching of muscles |
| 5 | Trachea filling | Trachea system becoming filled with air |
| 6 | Hatching | Larvae exits from the embryonic egg shell |

### Statistics

From our experience with spatial patterning and also from observing that larvae often hatch at similar times, we expect the relative temporal error (coefficient of variation, *CV*_τ_ = s.d. / mean) to be small (~2-5%), with a similarly small standard deviation (around 1-2%). We performed a power analysis to estimate the required minimum number of embryos for each condition. To observe a difference between a mean *CV*_τ_ = 0.04±0.02 and *CV*_τ_ =0.05±0.02 with power 0.8 requires n=34 samples (calculated using t-test). P-values were calculated (unless otherwise stated) using a two-tailed t-test comparison. In all datasets for *D. melanogaster, D. simulans* and *D. virilis*, n>35. Error on the temporal variation was estimated using Bootstrapping, with 100 simulations performed per data set.

### Modelling

The simulations were performed in Matlab. The experimentally measured time between landmarks at each temperature was used to determine the corresponding values of *λ*, the input parameter for the Gaussian distribution (with standard deviation *λ*^1/2^). We use a Gaussian distribution rather than the Erlang distribution (which describes the distribution of time between events in a Poisson process) since we lack sufficient information to reliably parameterise the Erlang distribution. The data for the first time point (corresponding to germband retraction) was distributed as measured experimentally, as there was no previous time course data available – as cephalic furrow is used to define time 0. For subsequent events, the history dependence was implemented as described in text. For each temperature, 1000 simulations were performed, where random numbers were generated using the Matlab function *randn*. Fitting of *r,* the history-dependent parameter, was done to data at 16°C, 21°C and 25°C for each species. We first attempted to use a single value for all three temperatures. For *D. simulans*, this gave a good fit to the data, but for *D. virilis* and *D. melanogaster* this resulted in a poor fit at least at one temperature. For *D. melanogaster, r=0* resulted in a poor fit to the data except at low temperatures (figure S5*a*). Likewise, using *r=40mins* or *r=-40mins* for all data resulted in a poor fit (figure S5*b* and figure S5*c*, respectively). Therefore, for *D. virilis* and *D. melanogaster* we allowed two values of *r*, depending on temperature, as outlined in the Results. Note, our approach with the model was not to find the best value of *r* at each temperature, but to find a minimal range of values for *r* that can explain as wide a portion of the data as possible. To model the temperature shift experiments, the value of *r* corresponded to the temperature of the system at each particular landmark. However, we find that for shifts from 26°C, a better fit was achieved with *r=0,* not *r<0* after the temperature shift (figure 5*d*). Finally, the actual time of temperature shift for each embryo is slightly different. The time leading up to and immediately after the temperature shift were both drawn as random Gaussian variables and the relative contribution of each weighted to ensure that the average time of development corresponded to experimental measurements. For *D. virilis* data for muscle twitching was excluded due to experimental error in determining the onset of such twitching (figure S2*g*).

## Results

### Temporal development of *Drosophila* embryos is robust across a wide temperature range

Developmental time in *D. melanogaster* increases markedly as temperature decreases (figure 2*a*, 18). However, the temporal variation, *σ_τ_*, shows a more complicated behaviour. If temporal variability in embryonic development is largely due to random variations then we expect 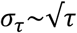. This corresponds to the temporal development of the embryo being a continuous process with little history dependence or checkpoints. However, we find that embryos developing in the range 19-21°C typically have reduced variation than at both high (>23°C) and low (<19°C) temperatures (figure 2*b*). At low temperatures, larger *σ_τ_* is unsurprising, due to accumulation of more error from the longer developmental time. The increased temporal error at higher temperatures is surprising given that developmental time is significantly shorter.

As the mean developmental time varies drastically across temperatures, we reason that a more appropriate measure is the coefficient of variation, *CV*_τ_ = *σ_τ_*/*τ*. This dimensionless measure enables comparison of developmental noise while accounting for changes in developmental time with temperature. This is analogous to quantification of the spatial precision of boundaries, where the boundary position is typically scaled by the embryo length. If the temporal variability is dominated by random noise, then we expect 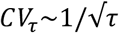. Remarkably, for all temperatures in the range 16°C-28°C, *CV*_τ_ was less than 6% for all landmarks except hatching (figure 2*c*). At very high temperatures *CV*_τ_ increases further, with larger embryo-to-embryo variability at 30°C (figure 1*c*). Indeed, embryos developing at 30°C have a temporal variation around 4-5 times larger than embryos developing at 21°C.

**Figure 2:**
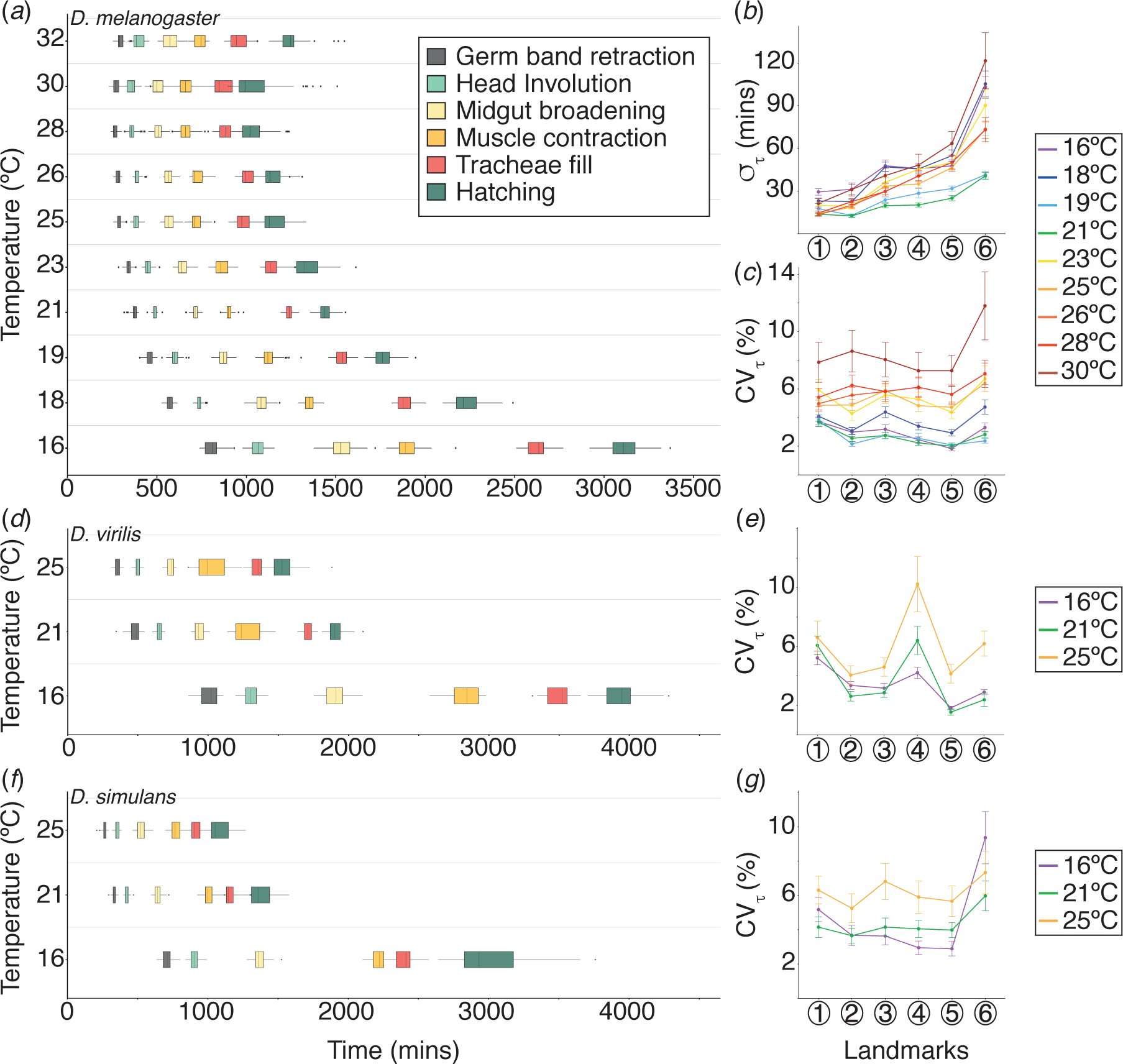
*Drosophila* embryonic development is temporally robust. (*a*) Distribution of developmental times at each landmark scored for *D. melanogaster*. (*b*) Absolute error in developmental time at each landmark for *D. melanogaster* at different temperatures. (*c*) Relative error in developmental time at each landmark for *D. melanogaster* at different temperatures. (*d*) Distribution of developmental times at each landmark scored for *D. simulans*. Colour coding as (*a*). (*e*) Relative error in developmental time at each landmark for *D. simulans* at 16, 21 and 25°C. (*f*) Distribution of developmental times at each landmark scored for *D. virilis*. Colour coding as (*a*). (*e*) Relative error in developmental time at each landmark for *D. virilis* at 16, 21 and 25°C. All error bars are standard deviation, which in (*c*,*e*,*g*) are estimated by bootstrapping.

For T<23°C, we observed a gradual decrease in *CV*_τ_ during development, qualitatively consistent with 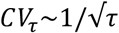. However, *CV*_τ_ at higher temperatures was significantly larger than at low temperatures (e.g. p=0.003, comparing mean *CV*_τ_ at midgut broadening for embryos developing in range 23-26°C with those below 21°C). Further, *CV*_τ_ was approximately constant throughout embryonic development for embryos developing above 23°C (figure 2*c*). Although temporal development is robust at high temperatures, there are clear differences in behaviour in *CV*_τ_ compared with lower temperatures.

We next explored the temporal robustness of development in two related *Drosophila* species, *D. virilis* and *D. simulans,* which diverged around 40MYA and 5MYA from *D. melanogaster* respectively (from flybase.org). We chose to focus on temperatures 16°C, 21°C, and 25°C as these represent the range of standard laboratory conditions for *Drosophila. D. virilis* develops in more temperate climates and has a significantly longer developmental time than *D. melanogaster* (figure 2*d*, S1*e* and Movie 1, 18). As with *D. melanogaster*, the absolute temporal error is higher at low temperatures (figure S1*f*). However, at high temperatures *CV*_τ_ is similar for *D. virilis* compared with *D. melanogaster*, with an average relative error of around 4.8% (figure 2*e*). Therefore, the temporal development of *D. virilis* is also robust across a wide range of temperatures.

*D. simulans* is a tropical species that develops faster than *D. melanogaster* (figure 2*f*, S1*e* and Movie 1, 18). *CV*_τ_ is around 4%, suggesting the *D. simulans* development is temporally robust. It is noteworthy though that *CV*_τ_ is larger in *D. simulans* than *D. melanogaster* at all landmarks (excluding hatching) for each temperature measured (p-value<10-2 for all conditions). Interestingly, for intermediate temperatures (T=21°C), *CV*_τ_ is relatively constant throughout development for *D. simulans*, in contrast to *D. melanogaster* where it decreases with developmental time. Again, we see that *CV*_τ_ at high temperatures is significantly greater than at low temperatures (p-value<10^−3^), even though embryo viability is similar (figure 2*g* and S1*c*).

As with *D. melanogaster*, both *D. virilis* and *D. simulans* have larger absolute temporal errors at 16°C (figure S1*f-g*). The absolute temporal error at 21°C and 25°C is surprisingly similar for both species. Finally, we tested more systematically the dependence of *CV*_τ_ on the developmental time. Fitting *CV*_τ_ = *aτ^s^* - for each species at each temperature, we find that *D. virilis* has relatively constant *s* around −0.5 at all temperatures tested (figure S1*h*). However, for both *D. melanogaster* and *D. simulans*, *s* approaches zero at higher temperatures. We return to these observations below when we explore the temporal correlations during development.

Overall, we see that all three species analysed are temporally robust except at very high temperatures. However, there is clear temperature dependence in the variability of the temporal trajectories of developing embryos and we explore this further below.

### Temporal coordination in varying temperature environments

To investigate further the temporal trajectories and how they depend on temperature we recorded *D. melanogaster* embryos developing in varying temperature environments. First, we considered shifts of temperature 16°C to/from 21°C (figure 3*a-b*) and 21°C to/from 26°C (figure S2) after head involution. We chose these temperatures such that the temperature change was ±5°C from 21°C. Clear shifts are observable in the timing error (figure 3*c*) but these appear largely transient. Looking at *CV*_τ_ we see that decreasing or increasing the temperature to or from 21°C after head involution results in acute increase of the relative temporal error (figure 3*d* and figure S2). Therefore, the temporal development of *D. melanogaster* is surprisingly robust to abrupt temperature variations.

**Figure 3:**
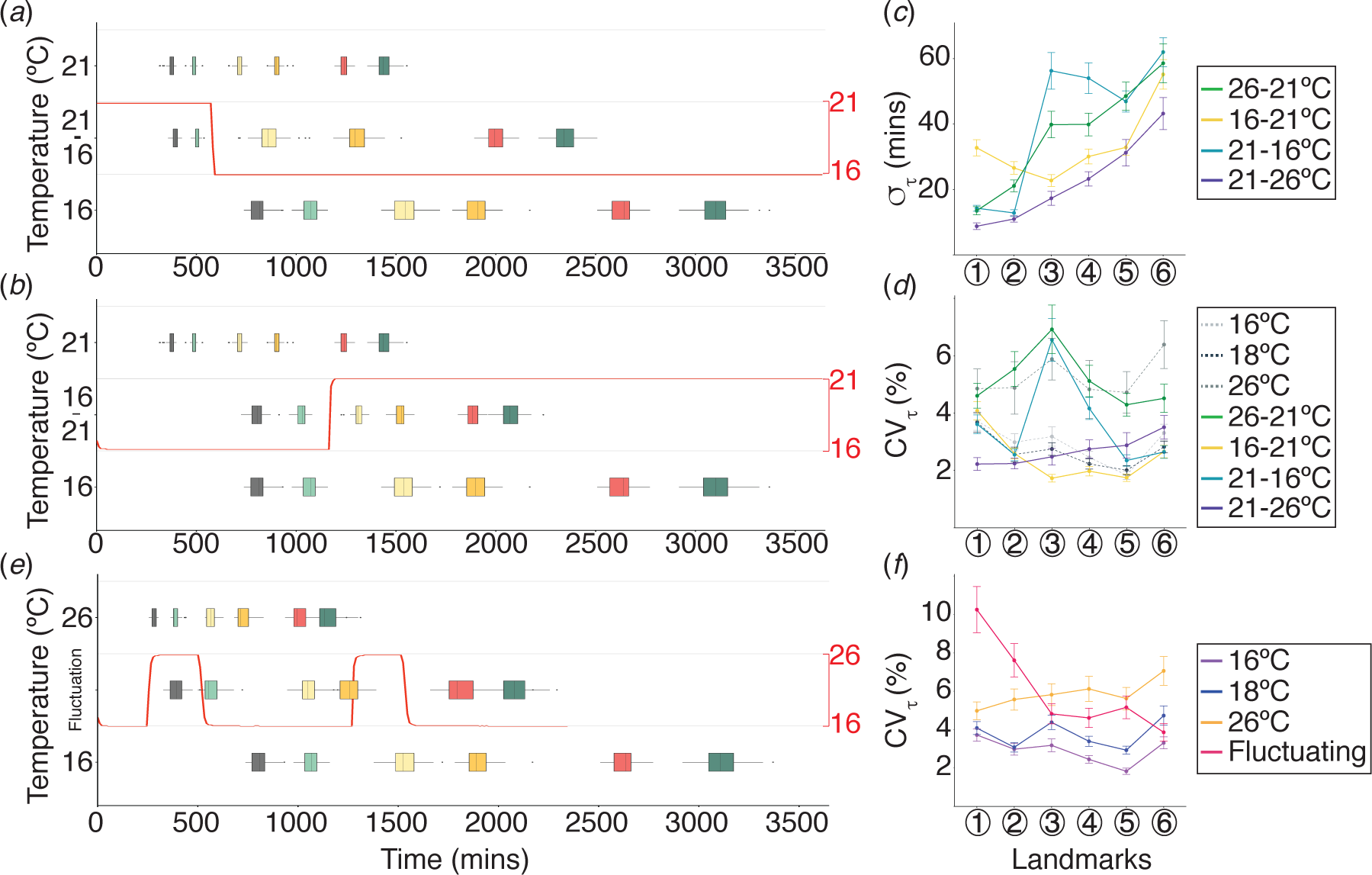
*D. melanogaster* embryonic development under thermal perturbations. (*a*,*b*) Distribution of developmental times at each landmark scored for *D. melanogaster* under 16 to/from 21°C temperature shift. Temperature was shifted after head involution and is represented by a red curve. Colour coding as in Figure 1a. (*c*) Absolute error in developmental time at each landmark for *D. melanogaster* in 16 to/from 21°C and 21 to/from 26°C temperature shift experiments. (*d*) Relative error in developmental time at each landmark for *D. melanogaster* in 16 to/from 21°C and 21 to/from 26°C temperature shift experiments. (*e*) Distribution of developmental times at each landmark scored for *D. melanogaster* in temperature pulse experiments. Temperature was shifted at regular interval and is represented by a red curve. Colour coding as in Figure 1a. (*f*) Relative error in developmental time at each landmark for *D. melanogaster* in pulse experiments. All error bars are standard deviation, which in (*d*,*f*) are estimated by bootstrapping.

To further test the temporal robustness of *D. melanogaster*, we recorded the temporal development of embryos at 16°C applied with two +10°C temperature pulses of 4-hour duration during development (figure 3*e*). This temperature range was chosen as it represents the regime of robust temporal development. Measuring *CV*_τ_, we see that the first temperature pulse results in a large temporal perturbation but this shift is largely negated by midgut broadening. The second pulse results in a much smaller shift in *CV*_τ_. After midgut broadening, the average *CV*_τ_ is between 26°C and 18°C (the overall average temperature throughout development) (figure 3*f*). This suggests that the temporal trajectories can partially – though not completely – adapt to abrupt shifts in temperature.

### The temporal coordination between developmental landmarks is temperature dependent

To better understand how such temporal robustness emerges, we investigated how the timing of developmental landmarks depended on the developmental history of the embryo. We performed a covariance analysis across all landmarks for *D. melanogaster* at constant temperatures to examine how correlated the timings of later landmarks were with earlier events (figure 4*a* and S3*a*). The covariance is significantly reduced at 21°C compared to both lower and higher temperatures (table 2). These results corroborated with individual time courses, which indicated that embryos tend to continue on the same temporal trajectory throughout development at low and high temperatures: e.g. an embryo that is developing (relatively) fast early on, also develops (relatively) faster later in development (figure 4*b*). The low covariance at intermediate temperatures is due to greater variability in the temporal trajectories.

**Figure 4:**
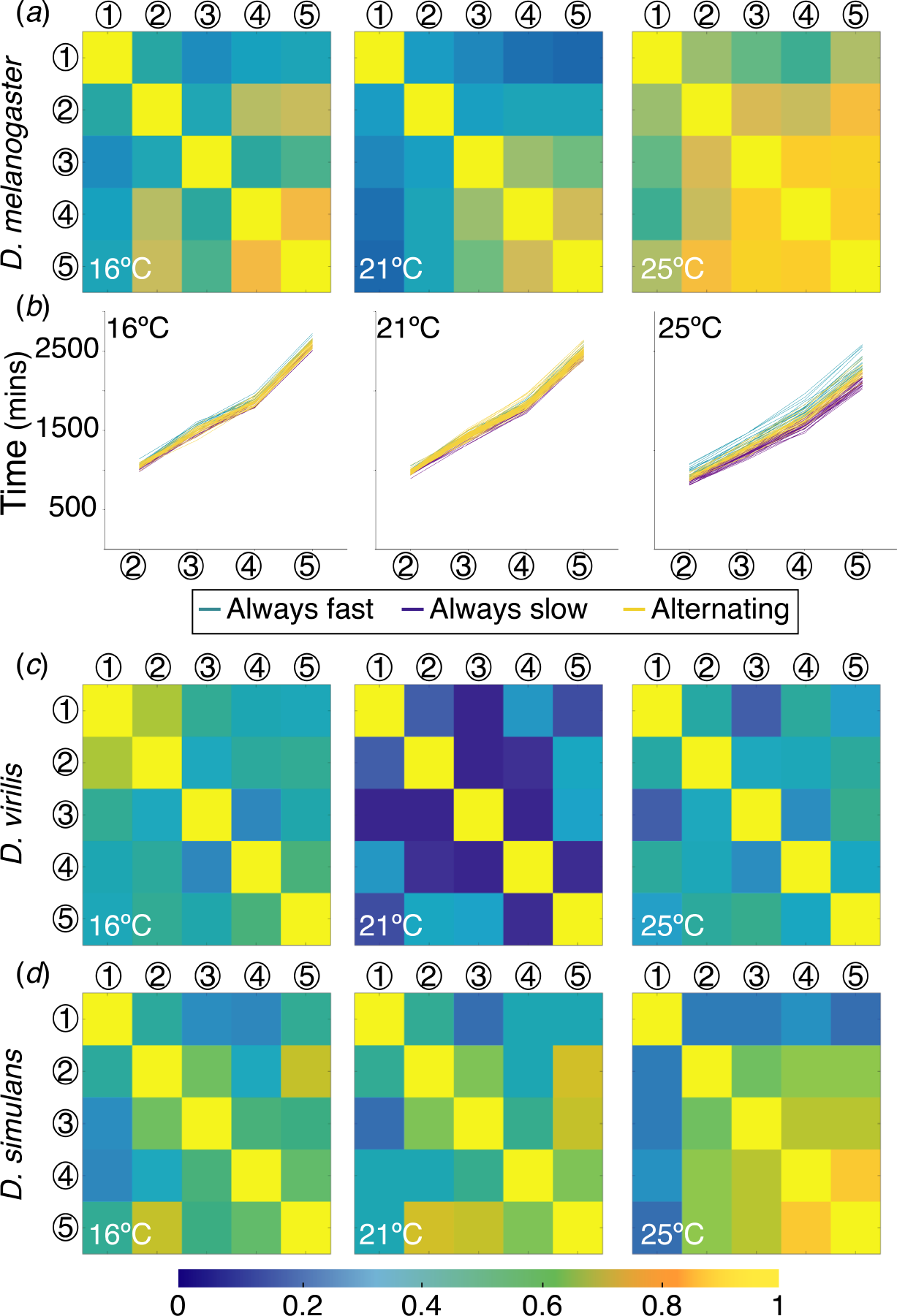
Temporal trajectories are history dependent. *(a)* Covariance between each landmark for *D. melanogaster* at 16, 21, and 25°C. (*b*) Individual embryo time courses at 16, 21 and 26°C colour coded by whether an embryo’s development is always faster than the mean at that temperature (turquoise), always slower than the mean at that temperature (purple), or whether the trajectory alternates at least once between being faster or slower than the mean (yellow). (*c*, *d*) Covariance between each landmark at 16, 21, and 25°C for *D. virilis* (*c*) and *D. simulans* (*d*).

**Table 2.** CV and covariance (cov) relative to landmark (1) in temporal trajectories in *D. melanogaster, D. simulans* and *D. virilis*.

| Species | Temperature<br>°C | Landmark |  |  |  |  |  |  |  |  |  |  |  | p<br>CV | p<br>Cov |
| --- | --- | --- | --- | --- | --- | --- | --- | --- | --- | --- | --- | --- | --- | --- | --- |
|  |  | 1 |  | 2 |  | 3 |  | 4 |  | 5 |  | 6 |  |  |  |
|  |  | CV% | Cov | CV<br>% | Cov | CV<br>% | Cov | CV<br>% | Cov | CV<br>% | Cov | CV<br>% | Cov |  |  |
| *D.*<br>*melanogaster* | 16 | 3.7 | n/a | 3.0 | 0.48 | 3.2 | 0.29 | 2.5 | 0.39 | 1.8 | 0.42 | 3.3 | 0.34 | 0.67 | 0.046 |
|  | 21 | 3.7 | n/a | 2.6 | 0.35 | 2.8 | 0.28 | 2.2 | 0.20 | 2.0 | 0.16 | 2.8 | 0.13 | n/a | n/a |
|  | 25 | 5.0 | n/a | 5.6 | 0.76 | 5.8 | 0.69 | 6.1 | 0.69 | 5.6 | 0.72 | 7.1 | 0.72 | <10<sup>-4</sup> | <10<sup>-4</sup> |
|  | 16-21 | 4.1 | n/a | 2.6 | 0.48 | 1.7 | 0.26 | 2.0 | 0.30 | 1.7 | 0.28 | 2.7 | 0.30 | 0.07 | 0.56 |
|  | 21-16 | 3.6 | n/a | 2.6 | 0.67 | 6.6 | 0.36 | 4.2 | 0.22 | 2.4 | 0.32 | 2.6 | 0.30 | 0.49 | 0.17 |
|  | 21-26 | 2.2 | n/a | 2.2 | 0.13 | 2.5 | 0.36 | 2.8 | 0.24 | 2.9 | 0.26 | 3.5 | 0.23 | 0.05 | <10<sup>-4</sup> |
|  | 26-21 | 4.6 | n/a | 5.5 | 0.79 | 6.9 | 0.73 | 5.1 | 0.71 | 4.3 | 0.78 | 4.5 | 0.80 | 0.75 | 2x10<sup>-4</sup> |
| *D.*<br>*simulans* | 16 | 5.2 | n/a | 3.7 | 0.54 | 3.6 | 0.32 | 3.0 | 0.31 | 2.9 | 0.55 | 9.4 | 0.47 | 0.08 | 0.89 |
|  | 21 | 4.2 | n/a | 3.7 | 0.55 | 4.2 | 0.24 | 4.1 | 0.49 | 4.0 | 0.49 | 6.0 | 0.46 | n/a | n/a |
|  | 25 | 6.3 | n/a | 5.3 | 0.27 | 6.8 | 0.27 | 5.9 | 0.67 | 5.7 | 0.25 | 7.3 | 0.23 | 0.04 | 0.065 |
| *D.*<br>*virilis* | 16 | 3.3 | n/a | 2.6 | 0.59 | 2.7 | 0.51 | 3.3* | 0.42 | 1.3 | 0.26 | 1.3 | 0.40 | 0.97 | 0.15 |
|  | 21 | 3.4 | n/a | 2.4 | 0.39 | 2.5 | -0.05 | 7.0* | 0.50 | 1.4 | 0.17 | 2.8 | 0.10 | n/a | n/a |
|  | 25 | 5.5 | n/a | 3.6 | 0.22 | 3.2 | -0.02 | 10* | 0.43 | 2.8 | 0.29 | 4.0 | 0.34 | 0.11 | 0.60 |

For experiments without temperature shift, the p-value is calculated from comparing the mean CV and Cov for landmarks 2-5 (i.e. excluding hatching), relative to 21°C. For temperature shift experiments, the mean CV and Cov are calculated for landmarks 4-5 (after temperature shift) and compared with the mean CV before the shift (landmarks 1 and 2) and Cov of embryos maintained at a constant temperature (at temperature after shift) respectively. \* large CV due to experimental error in identifying muscle twitching in *D. virilis*, not biological variation.

We also calculated the covariance in developmental times across different landmarks for *D. virilis* (figure 2*c*) and *D. simulans* (figure 2*d*). For *D. virilis,* a similar trend was observed as for *D. melanogaster* (table 2). However, it is noteworthy that the covariance at high temperatures is significantly less than that in *D. melanogaster.* In contrast, the covariance of *D. simulans* was very similar to that of *D. melanogaster,* except for correlations with germband retraction at 25°C.

To test this last observation further, we checked how the proportion of temporal trajectories that were always fast or slow changed if we excluded germband retraction (figure S3*b*). For *D. virilis*, little change was observed with 74% and 72% of the temporal trajectories varying about the mean developmental times when beginning from and after cephalic furrow respectively at 25°C. For *D. simulans* there was a larger change in the proportion of track types with mixed trajectories at 25°C (62% to 49%, p = 0.25) but the shift itself was not significant. However, comparing *D. virilis* and *D. simulans*, we see that excluding germband retraction resulted in a significant difference between the proportion of embryos with mixed temporal trajectories in *D. simulans* and *D. virilis* (p=0.03). For *D. melanogaster* there was a marked decrease in the proportion of track types with mixed trajectories at 25°C (43% to 27%, p=0.01). These results suggest that the temporal development of *D. melanogaster* and *D. simulans* are more sensitive to temporal variability early in embryogenesis, though further experiments, particularly on *D. simulans*, will be needed to confirm this observation.

We also performed the covariance analysis for the temperature shift experiments. The shift from 26°C to 21°C resulted in continuing high covariance between developmental landmarks, which explains the large *CV*_τ_ despite reduction to 21°C (figure S3*c*). In contrast, embryos initially raised at 21°C but then shifted to 26°C retained a relatively small covariance (figure S3*c*). For changes to and from 16°C, the covariance behaviour was similar (figure S3*c*). Therefore, we see that development at a particular temperature during early development affects the heterochronicity of later processes.

### A single correlation parameter can explain the temperature dependence of *CV*_τ_

To better understand these observations, we simulated developmental temporal trajectories. In each simulation, we have six landmarks, occurring at times *τ_i_*, with the time between each landmark and the distribution in times for the first landmark defined from the experimental data. The developmental time for subsequent landmarks is calculated as 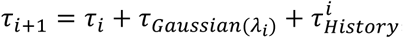, where *τ*_*Gaussian*_(*λ_i_*)__ is drawn from a Gaussian distribution with mean *λ_i_* and standard deviation 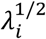, determined by the experimentally measured time between landmarks *i* and 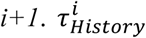 represents correlations between the timing of previous landmarks and the subsequent landmark. For simplicity we take the form 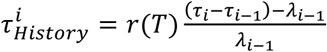, where *r* is a (temperature-dependent) constant. If *r=0*, there is no history dependence. For *r*<0, temporal variations are reduced by, for example, slowing down temporal trajectories that are faster than the mean population development time. For *r*>0, temporal trajectories that are faster than the mean population development time are reinforced, increasing the temporal error.

This simple model for embryonic temporal development fits the observed *CV*_τ_ with a temperature dependent *r_mel_* where *r_mel_ (T*>23°C) ≈+*40 mins*, *r_mel_ (T*<23°C) ≈−*40 mins* (figure 5*a*). Fluctuations at high temperatures are dominated by *τ_History_* whereas at low temperatures *τ_Gaussian_* dominates. For *D. virilis*, we predicted that since development is longer, there is greater potential for feedback to regulate developmental time. Consistent with this, we found *r_vir_(T*>23°C) ≈ *-25 mins*, *r_vir_ (T*<23°C) ≈ *-70 mins* for the *D. virilis* data (figure 5*b*). For *D. simulans*, we had the opposite prediction: since developmental time is faster, we expected that any history-dependence would amplify, rather than reduce, the temporal variability. Intriguingly, we found that a single parameter *r_sim_ = +35 mins* was able to fit our model the *D. simulans* data (figure 5*c*). Therefore, the different temporal trajectories of the three species can be encapsulated within a single parameter that defines the level of history-dependence and whether such history-dependence dampens (*r<0*) or amplifies (*r>0*) temporal fluctuations. For *D. melanogaster* the sign of *r* changes sign with temperature but for the faster developing *D. simulans r* is positive and for the slower developing *D. virilis r* is negative at all temperatures analysed.

**Figure 5:**
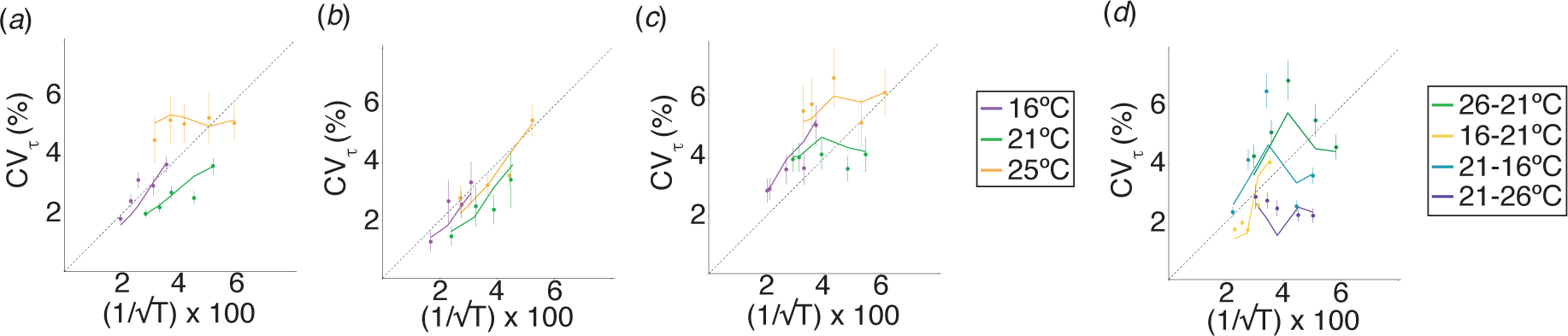
A single parameter defines the level of history-dependence. (*a*-*d*) Relative temporal error against 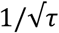 compared with model predictions: (*a*) *D. melanogaster, (b) D. virilis, (c) D. simulans* and (*d*) *D. melanogaster* with shifted temperature. Dashed line corresponds to 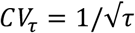. All error bars are standard deviation, estimated by Bootstrapping.

Finally, to further test our phenomenological model for the temporal trajectories we examined the temperature shift experiments. The model can qualitatively replicate our experimental observations using *r_mel_* as above, but imposing that *r_mel_=0* after the temperature shift from 26°C to 21° (figures 5*d*). In conclusion, we can qualitatively explain the observed behaviour of *CV*_τ_ at both high and low temperatures in three species with a single fitting parameter.

## Discussion and conclusion

Our results show that *Drosophila* temporal development is highly robust across three different species and a wide range of temperatures. Interestingly, this level of precision is similar to that of embryonic spatial precision of gene expression boundaries along the anterior-posterior axis in *Drosophila* (2). We find that the behaviour of the temporal trajectories is non-trivial, in the sense that the statistical correlations vary both with temperature and between species. It is notable that the tropical species have generally similar correlative behaviour, whereas the temperate *D. virilis* has distinct covariance.

At high temperatures in *D. melanogaster,* our simple model can replicate the observed variability with behaviour akin to reinforcement in the temporal trajectories, whereby trajectories that are fast (slow) compared to the mean developmental time are favoured to remain fast (slow) throughout development. At intermediate temperatures, we find behaviour akin to negative feedback, whereby trajectories that are fast (slow) compared to the mean developmental time are unlikely to remain fast (slow) throughout development. This is suggestive of temperature specific regulation. In contrast, (i) the more rapidly developing *D. simulans* has behaviour consistent with positive feedback at all temperatures analysed; and (ii) the more slowly developing *D. virilis* has behaviour consistent with negative feedback at all temperatures analysed. For rapidly developing embryos, a cohort of eggs laid at similar times will hatch close together even with noise in their temporal trajectories. Taking these observations into account, we reasoned that there may be some constraint on the *absolute* error – *i.e.* processes may exist to maximise the number of embryos within a cohort that hatch within a particular time window. Therefore, we went back and compared the absolute temporal errors between species. At 25°C all three species had very similar absolute error, *i.e.* the negative feedback in *D. virilis* at 25°C is sufficient to compensate for the longer developmental times (Figure S5). Conversely, at 16°C, *D. virilis* typically had larger absolute temporal error than the other species except near hatching. Note this is because the other two species have increased their temporal precision, not because of a decrease in *D. virilis* precision. These results suggest that regulatory mechanisms may exist to control developmental time, and these are tuned to respond to temperature variations. Interestingly, *D. melanogaster* shows high thermal tolerance in the embryo which is lost in the adult, suggesting that the embryo has active mechanisms to adjust for temperature changes (22) and these may play a role in regulating developmental time.

The assay presented here is a viable platform for understanding how embryos adapt to subtle changes in environment. Past investigations have generally focused on the effect of temperature at larval and adult stages. The differences in embryonic timing between species of *Drosophila* has been quantified and shown to obey the Arhennius rule for reaction rates (18). Recent work has shown that *Drosophila* raised at a specific temperature did not select for flies optimal (in terms of fecundity) at that temperature (23). Rather, flies exposed to temporally varying temperatures displayed increased fecundity across a broad temperature range, outperforming flies maintained at specific temperatures. Temporally variable environments have also been shown to delay reproductive maturation (24,25). The evolution of adaptability to temperature changes has also been intensively studied (8). Our study addresses an important gap; understanding the effect of temperature on an immobile, and therefore vulnerable, entity. Evolution has shaped *Drosophila* embryos to be able to cope with a wide range of temperatures. Strikingly, acute temperature changes across the natural physiological range has a modest effect on the temporal variability. Essentially, our results indicate that *Drosophila* embryos have the machinery in place to adapt to temperature changes.

We noticed interesting differences in the behaviour of tropical and temperate species. To test this observation further, we obtained the original dataset from (18) which covered 11 different *Drosophila* species. The number of embryos for each species and temperature ranged from 6 to over 70. These embryos were collected in a different environment and under different imaging conditions (e.g. we did not dechorionate the embryos) making direct comparison with our results difficult (Figure S6*b*). Due to the variable sample sizes, we consider two sets: (i) tropical (*D. simulans, D. ananassae, D. sechellia, D. willistoni, D. yakuba, D. erecta*); and (ii) non-tropical (*D. virilis, D. mojavansis, D. persimilis, D. pseudoobscura*). Comparing the change in *CV*_τ_ between cellularisation and trachea filling, we find that tropical species show significantly more variability in *CV*_τ_ as temperature is varied than non-tropical species (figure S6*c-d*). This is consistent with our above results and suggests that species that are exposed to wider temperature fluctuations have developed regulatory processes to buffer the effects of such temperature changes on developmental time. However, more detailed species-specific analysis will be required to confirm this observation. Along these lines, a recent observation revealed that the *Drosophila* βtubulin97EF is upregulated at low temperatures and contribute to stabilize microtubules (26). This example demonstrates that differential regulation of intracellular components is necessary for acclimation to environmental changes. Further, miRNAs and small non-coding molecules may play a role in temporal regulation as these molecules are known to buffer noise and to respond to environmental changes (27,28).

The timing noise in single cells has been quantified in a number of systems (29,30). Work on bacterial cells has shown that changes in temperature alter both the cell growth rate and the time to division equally (31) and this can be explained within a cyclic autocatalytic reaction whereby each element catalyses the next element (32). In particular, temperature-dependent scaling of cellular time is sufficient to explain experimental observations of bacterial growth response to temperature changes. It will be interesting to extend our simple model to see whether such general physical principles also apply to developing systems.

The careful control of timing during development, for example in the segmentation clock (33,34), and in adults, such as circadian rhythms (35), is essential for life. Yet, heterochronicity has been shown to be crucial for the evolution of new traits. By altering the timing or sequence of developmental effects, new features can emerge, such as increased segment number in snakes (36,37). Our quantitative results demonstrating a temperature specific response in the temporal trajectories of development are suggestive of factors regulating the timing of development. In the larvae, a number of hormonal signals have been identified that regulate developmental time (38,39), but currently little is known about mechanisms of temporal regulation in the embryo. Finally, there has been significant work in trying to understand ecological adaptation to changing environments (40), and it will be interesting to quantify the temporal trajectories of development in a continuously fluctuating environment.

## Author Contributions

J.C., C.A., and T.E.S. designed the experiments. J.C. performed the experiments and quantified developmental times. C.A. and T.E.S. analysed the data. T.E.S. performed the simulations. All authors contributed to writing of the paper.

## Acknowledgements

We thank Patrice Koehl for advice on the statistical analysis. T.E.S. was funded through a National Research Foundation Singapore Fellowship (NRF2012NRF-NRFF001-094).

